# Genomic mapping of the modifiers of *teosinte crossing barrier 1* (*Tcb1*)

**DOI:** 10.1101/2022.07.18.500501

**Authors:** Namrata Maharjan, Merritt Khaipho-Burch, Prameela Awale, Abiskar Gyawali, Vivek Shrestha, Yajun Wu, Donald L. Auger

## Abstract

Pollen cross-contamination has been a major problem for maize breeders. Mechanical methods applied to avoid cross-contamination are largely ineffective and time-consuming. Cross incompatibility barriers are genetic factors involved in maize fertilization that can be used as an effective method to prevent pollen cross-contamination. *Teosinte crossing barrier 1 (Tcb1)* is a cross-incompatibility system in which silks possessing dominant *Tcb1-s* reject pollen possessing the recessive allele *(tcb1)*. However, successful fertilization occurs when *Tcb1-s* pollen falls upon *tcb1* silks or under self-fertilization of *Tcb1-s* pollen on *Tcb1-s* silks. Previous studies have shown that the efficacy of dominant *Tcb1-s* was reduced when repeatedly backcrossing with maize inbred lines suggesting the presence of modifiers to *Tcb1-s*. To find those modifiers, we conducted a QTL mapping experiment using the Intermated B73 x Mo17 (IBM) recombinant inbred lines (RILs) for two consecutive years. Two significant and stable QTL were identified on chromosomes 4L and 5S explained 16% and 17.6% of the total phenotypic variation (R^2^), and both had negative additive effects. Further investigation of these QTL regions identified twelve candidate genes that could modify *Tcb1-s* activity. The introgression of the *Tcb1-s* genetic system, and its appropriate modifying factors, could be a novel and reliable solution for cultivar isolation in maize breeding.

## Introduction

Maize is a highly cross-pollinated crop. Pollen from one field can easily travel and pollinate silks in nearby fields, thus contaminating maize cultivars. Pollen contamination is a serious problem among farmers and breeders working with organic maize, landraces, sweetcorn, and inbred lines, leading to the loss of genetic purity. Different mechanical measures like increasing isolation distance and managing different planting dates have been in practice to avoid unwanted pollen from a neighboring field. For instance, displacing planting times by four to five days led to a 25 % reduction in the cross-fertilization rate, whereas a 6 to 10-day shift in planting date led to a 50-70% reduction in cross-contamination (Della Porta et al. 2008, Kozjak et al. 2011). However, these methods are not always practical to implement and are not completely reliable. Several genetic systems have been identified in the maize that avoids foreign pollen during reproduction. Cross-incompatibility (CI) is a pre-zygotic fertilization barrier that prevents pollen from a foreign pollen parent from successfully fertilizing an egg. Maize has nine known cross-incompatibility systems located throughout the genome, *gametophyte factor 1* (*ga1*, 4S), *ga2* (5L), *ga3* (7), *ga4* (1), *ga6* (1), *ga7* (3L), *ga8* (9S), *ga10* (5S), and *teosinte crossing barrier 1* (*tcb1*, 4S) (Burch 2018, Nelson 1994, Neuffer et al. 1997).

*Teosinte crossing barrier 1* is one type of unilateral cross-incompatibility factor that exhibits a selective preference for its own genotype and has potential uses in maize breeding programs (Kermicle and Evans 2010, Kermicle et al. 2006, Lu et al. 2014). *Tcb1* was first described by Kermicle and Allen (1990) in *Zea mays* ssp. *mexicana* collected from Central and Southern Mexico. Initially, it was named the Teosinte Incompatibility Complex (TIC). Two accessions of *Zea mays spp. mexicana* from the Central Plateau and Chalco were introgressed into a W22 inbred background to characterize the TIC locus further. This locus harbor two tightly linked genes: *Tcb1-m (male)* and *Tcb1-f (female)* (Burch 2018, Kermicle 2006, Lu, Kermicle and Evans 2014). Depending upon the presence and absence of these two genes, the *Tcb1* locus has four haplotypes. These haplotypes have only the male function, only the female function, have both male and female functions represented as *Tcb1-s (strong)*, and lacking both functions represented as *tcb1* (Kermicle 2006, Lu, Kermicle and Evans 2014). This gene locus was mapped to the short arm of chromosome 4 and is 6 centiMorgans (cM) distal to *sugary-1* and 44 cM proximal to *ga1* (Evans and Kermicle 2001) (Figure 1).

**Figure 1:**
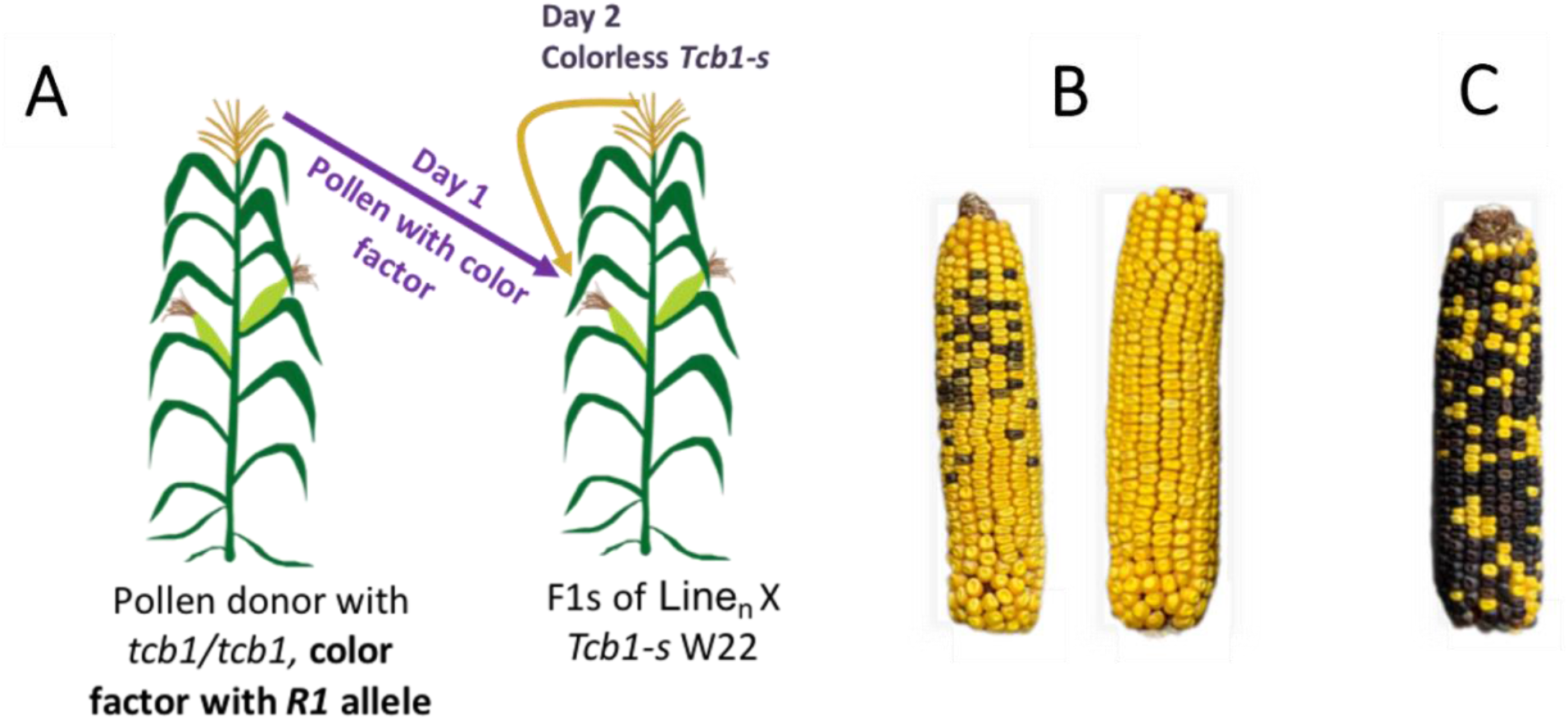
Visual methods to test of the strength of *Tcb1-s* efficacy in the F1 IBM RIls (*tcb1*/*Tcb1-s*). A) The efficacy of *Tcb1-s* in the F1 lines was evaluated by pollinating with *R1 C1 tcb1* pollen the first day and on the second day the plant was self-pollinated. B) Strong efficacy of *Tcb1-s* is indicated if the resulting ear has few or no blue kernels. (C) A weak effect of *Tcb1-s* is indicated if the ear being heavily contaminated with blue kernels.

The two best-characterized cross-incompatibility systems in maize are *ga1* and *tcb1*. In both cases, the cross-incompatibility systems act similarily (Burch 2018, Shrestha 2016). When the recessive allele (i.e., *ga1, tcb1*) is present in the pistil, it can accept pollen containing any other allele. When a strong dominant allele (i.e., *Ga1-s, Tcb1-s*) is present in the pistil, pollen containing the recessive allele is rejected, while pollen possessing the strong allele or the ‘male-type’ allele (i.e., *Ga1-m, Tcb1-m*) are accepted. The male-type alleles (i.e., *Ga1-m, Tcb1-m*) in the pollen grain can overcome the *Tcb1-s* barrier and are compatible with the strong-type allele (i.e., *Ga1-s, Tcb1-s*). However, plants with the male-type alleles can be fertilized by any other pollen alleles types (Nelson 1994). Both *Ga1-s* and *Tcb1-s* have been patented (Hoegemeyer 2005, Kermicle, Evans and Gerrish 2006).

Pollen of recessive *tcb1* cannot pollinate silks possessing dominant *Tcb1-s*, resulting in unsuccessful fertilization (Kermicle 2006, Kermicle and Evans 2010, Lu, Kermicle and Evans 2014). The molecular mechanisms behind this unsuccessful fertilization have shown a disruption of *tcb1* pollen tube growth in the pistil through pectin methylesterase activity encoded by *Tcb1-s* (Lu et al. 2019). The growth of *tcb1* pollen tubes was terminated at the tip of *Tcb1-s* pistils. *tcb1* pollen can only travel through 28% of the silk’s distance (LU *et al*. 2019). The thick cluster of callose was layered on the cell wall of the *tcb1* pollen tube tip in *Tcb1-s* silks (Lu, Kermicle and Evans 2014), resulting in the arrest of pollen tube growth. The thickening of callose had been observed in other self-incompatible crosses where pollen tube growth was halted (Geitmann et al. 1995). RNA-seq analysis of *Tcb1-s* revealed that *Tcb1-s* pistils had higher gene expression of TCB1-F/PME38, a homolog of pectin methylesterases 38 (PME38) belonging to group 1 type PME (Lu, Kermicle and Evans 2014). This protein removes methyl from methyl-esterified pectins from the pollen tube cell wall making it stiff, thus disrupting the pollen tube growth (Lu, Hokin, Kermicle, Hartwig and Evans 2019).

There is evidence that *Tcb1-s* does not act alone (Burch 2018, Evans and Kermicle 2001, Lu, Kermicle and Evans 2014). In addition to a separable *Tcb1-m* allele, additional modifiers influence *Tcb1-s* activity. A full-strength *Tcb1-s* containing positive modifiers can block *Ga1-s* and *ga1* pollen with only a 3% receptivity rate. An attenuated version of *Tcb1-s* lacking modifiers accepts more *Ga1-s* and *ga1* pollen with a 10% receptivity rate(Evans and Kermicle 2001). When backcrossed into W22 for ten generations, *Tcb1-s* activity decreased over time(Lu, Kermicle and Evans 2014). The progressive loss of *Tcb1-s* activity was suggested to be caused by epigenetic silencing of the *Tcb1-s* allele and potentially the segregation of modifying loci. Evidence for the presence of modifying loci of *Tcb1-s* between the inbred lines B73 and Mo17 has been observed (Kermicle, personal communication). This hypothesis was tested in Burch (2018) using the Intermated B73 Mo17 recombinant inbred line population (IBM RILs) crossed with a homozygous W22 *Tcb1-s/Tcb1-s* inbred line. The resulting F1s were polymorphic in *Tcb1-s* activity. Mapping of *Tcb1-s* activity within the IBM RILs showed evidence that *Tcb1-s* does not work alone and requires the action of modifiers to confer a stronger pistil barrier (Burch 2018). However, consistent and robust QTL regions were not identified due to a small sample size (77 RILs) and unreplicated trials.

In this study, we used the entire IBM population within replicated trials crossed with a homozygous W22 *Tcb1-s/Tcb1-s* to uncover consistent modifiers of the *Tcb1-s* locus within our F1 population. We performed QTL mapping for two consecutive years to identify and narrow down modifier QTL intervals and highlight twelve candidate genes that could modify the *Tcb1-s* cross-incompatibility mechanism. Understanding how the *tcb1* mechanism behaves in different genetic backgrounds is critical to their application in isolating commercial varieties during seed production. Our analysis revealed potential QTL that could be beneficial for marker-assisted introgression of *Tcb1* into commercial maize lines to restrict foreign pollen contamination.

## Materials and Methods

### Plant materials and generation of *Tcb1-s/tcb1* F1s in the IBM RIL population

302 recombinant inbred lines (RILs) from the intermated B73/Mo17 (IBM) population(Lee et al. 2002) (**Figure 4**) were collected from the Maize Genetic Cooperation Stock Center and used in this study. The population was planted in an experimental field at South Dakota State University in the summer of 2018 and 2019. The IBM RILs lack a functional *Tcb1-s* locus and instead carry the non-functional *tcb1* locus. To make F1s of IBM RILs, we crossed the population with homozygous *Tcb1-s* W22 stock and generated F1s (*Tcb1/tcb1*). The homozygous *Tcb1-s* stock was obtained from Dr. Jerry Kermicle from the University of Wisconsin Madison. Out of 302 IBM RILs, we were able to generate and plant 202 F1s in 2018 and 218 F1s in 2019. The planting rows were approximately 3 m in length, and plants were spaced about 23 cm apart (13 plants/row). In addition, we created and tested F1s with the founder lines B73 and Mo17 to create heterozygous B73 *tcb1*/*Tcb1-s* and Mo17 *tcb1*/*Tcb1-s* for both years.

### Efficacy testing of generated *Tcb1-s/tcb1* F1s

The methodology of fertilization efficacy testing was previously described (Burch 2018, Shrestha 2016). Briefly the efficacy of *Tcb1-s* in the F1 lines was evaluated by pollinating the F1 silks with *R1* (*colored 1*) *C1* (*colored aleurone 1*) *tcb1* pollen on the first day, and on the second day the plant was self-pollinated **(Figure 1A)**. Five F1 *Tcb1-s/tcb1* individuals were tested for each IBM RIL. None of the IBM *tcb1*/*Tcb1-s* F1 lines contained the alleles required for aleurone anthocyanin color (i.e., they were homozygous *r1, c1*, or both). 202 F1s generated in 2018 were used for efficacy testing in the summer of 2019, while 218 F1s generated in 2019 were used for efficacy testing in 2020. We used a stock containing *R1* and *C1* color factors in 2019. Due to stock availability, in 2020, a different stock was used that instead contained *R1-sc:m2* (an allele of *R1*). The testing stocks may have a slight discrepancy in kernel color. However, both testers possess the homozygous *tcb1* allele and produce enough of their respective color factors to distinguish between colored and non-colored kernels while testing the efficacy of generated F1s. Several rows of colored *tcb1* pollen donors were planted over four weeks in10 days intervals to provide a consistent supply of pollen during the time that F1 RILs flowered. The efficacy of pollen blocking in each F1 line was evaluated as described previously **(Figure 1B, C)**. Strong efficacy of *Tcb1-s* was indicated if the resulting ear had few or no blue kernels **(Figure 1B)**, and a weak effect was indicated the ear being heavily contaminated with blue kernels **(Figure 1C)**. We obtained 196 and 202 F1s (*Tcb1-s/tcb1)* IBM RILs in 2019 and 2020, respectively (**Supplemental Table S1 and S2)**.

### Phenotyping

The five double-pollinated ears were harvested from each F1s (*Tcb1-s/tcb1)* IBM RIL line. The ears were scored for the number of anthocyanin-colored kernels in both years. In 2019, we used the standard scale developed in our lab (**Figure 3**), where “0” indicated no colored kernels, “1” indicated up to 4% colored kernels, “2” for up to 8% colored kernels, “3” for up to 16% colored kernels, “4” for up to 32% colored kernels, “5” for up to 32% colored kernels and “6’’ for up to 64% and exceeding colored kernels **(Figure 2)**. This scale was developed to show a two-fold change in the number of contaminated color kernels. The mean efficacy score for each F1 ear was calculated and used as phenotype data for QTL analysis for 2019. In 2020, we did not use a standardized scoring scale because most ears had few colored kernels, so counting was more applicable than using a visual scale for scarcely contaminated ears. We counted the number of colored kernels and calculated the percentage of colored kernels present as a score for the ears. The mean percentage for all ears was calculated and used as phenotype data in 2020.

**Figure 2:**
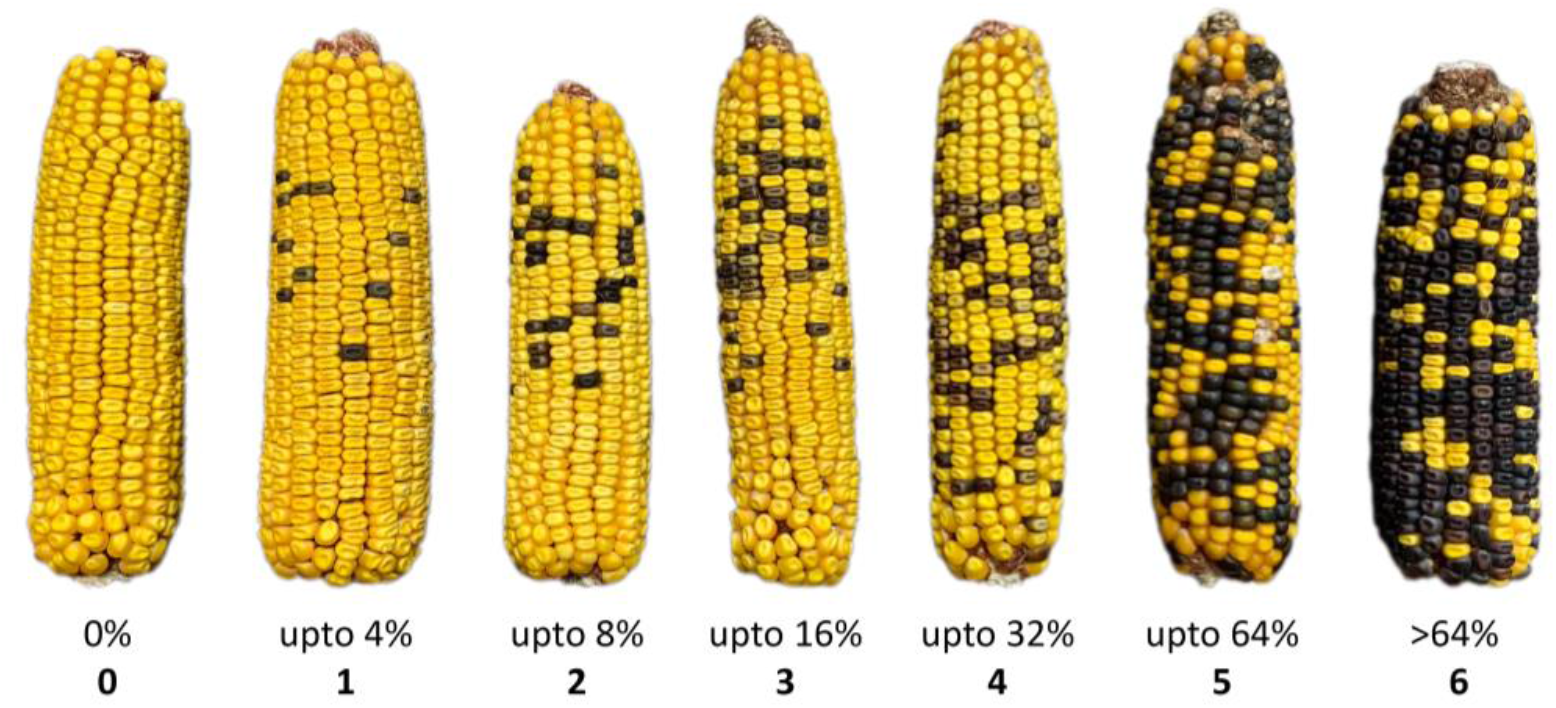
Phenotyping using the standardized scale system. Phenotyping of the matured ears using standardized scale: 0 = no colored kernels, 1 = up to 4% colored kernels, 2 = up to 8% colored kernels, 3 = up to 16% colored kernels, 4 = up to 32% colored kernels, 5 = up to 64% colored kernels and 6 = more than 64% colored kernels.

**Figure 3:**
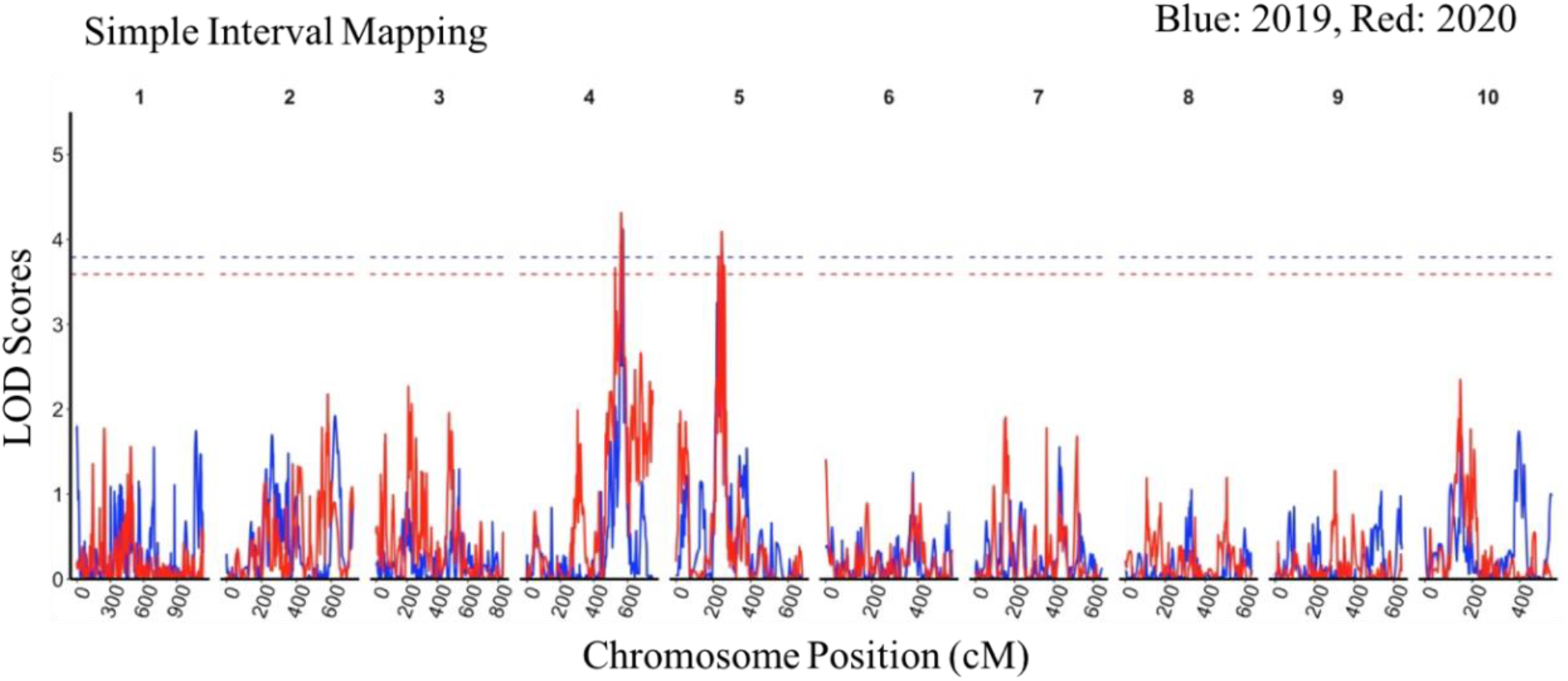
QTL mapping of modifiers of *Tcb1* in 2019 and 2020. The lower x-axis represents the chromosomal positon in centimorgan (cM) while the upper x-axis indicates the chromosome numbers. The y-axis represents the LOD scores. The vertical blue vertical lines represent the QTL result from 2019 whereas the red vertical lines represent the QTL results from 2020. The horizontal blue dotted line indicates 1000 permutated LOD threshold value of 3.70 for 2019 while the horizontal red dotted line indicates 1000 permutated LOD threshold value of 3.66 for 2020. The QTL map depicts significant and overlapping QTL peaks at chromosome 4L and 5S. Blue lines show 2019 results and red lines show 2020 results.

### QTL and candidate gene analysis

Genotype data for all the IBM RILs were obtained from MaizeGDB (https://www.maizegdb.org/mapscore_ibm302score) and used for QTL analysis. The average color kernel contamination was associated with markers by QTL mapping and analysis. The data was tested for normality (p-value <0.01, Shapiro-Wilk test) and normalized using quantile normalization in R. Two mapping approaches, simple interval mapping (SIM) and composite interval mapping (CIM), were used for this analysis. Both mapping approaches were carried out in R using the R/qtl package(Broman and Sen 2009).We used the Kosambi map function to estimate the map distance for all markers. We used Extended Haley Knott regression to conduct a chromosomal-wide one-way QTL scan at the interval of 1 cM. The significant LOD threshold was determined using 1000 permutation tests at an alpha level of 0.05. The significant QTL peaks were further explored to find the nearest significant marker. To estimate the confidence interval of QTL locations, a 1.5-LOD support interval was used. A two-dimensional, two-way QTL genome scan was conducted to detect linked loci and epistasis interaction that may present between the loci.

The genes located in the QTL intervals were explored for identifying candidate genes using MaizeGDB (B73_RefGen v3 gene model). The list of protein-coding genes was derived after excluding transposons, low confidence, uncharacterized, and unannotated genes. From this list, candidate genes were identified based on the protein families and functions affecting pollen, pollen development, pollen tube growth, and sterility.

## Results

### Phenotypic analysis of *Tcb1* efficacy in the founder lines (B73 and Mo17) and IBM RILs population

We evaluated the contamination scores from the F1s of founder lines (Mo17 (*tcb1*) × *Tcb1-s*) and (B73 (*tcb1*) × *Tcb1-s*). In 2019, the mean color contamination on the ears of the F1 Mo17 × *Tcb1-s* was 2.8% while for F1s of (B73 (*tcb1*) × *Tcb1-s*) was 3.6% **(Supplemental Figure 1A)**. In 2020, the mean contamination of of the F1 of (Mo17 (*tcb1*) × *Tcb1-s*) was 0% while for F1s of (B73 × *Tcb1-s*) was 2.7% **(Supplemental Figure 1B)**.

Across the F1s of the IBM RILs (tcb1) x *Tcb1-*s, we evaluated the contamination score within the phenotyped ears. The data used for this phenotypic characterization can be found in **(Supplemental Table 1,2)**. The mean contamination score in 2019 was 3.27 with a standard deviation of 1.22, while the mean contamination score in 2020 was 2.24 with a standard deviation of 1.45. The distribution of mean color contamination in the mapping population in both years deviated from normality **(Supplemental Figure 2A,C)**, and was normalized **(Supplemental Figure 2B,D)** using quantile normalization before QTL analysis.

### QTL analysis revealed significant and stable QTL on 4L and 5S

The simple interval mapping (SIM) analysis for the *Tcb1-s* modifiers identified two significant QTL on chromosomes 4L and 5S, and these results were consistent for both years (2019 and 2020) **(Figure 3, Supplemental Figure 3,4)**. The overall phenotypic variation (R^2^) explained by QTL was 16% and 17.6% in 2019 and 2020, respectively **(Table 1)**. We also used composite interval mapping (CIM) to perform the QTL analysis. Although CIM did not identify QTL that surpassed the permutated threshold, the results were very similar and followed the same QTL trends as SIM **(Supplemental Figure 5)**.

**Table 1:** Summary of significant QTL for 2019 and 2020. The table shows the year experiment was conducted, the QTL name, chromosome (Chr.), QTL peak position in cM, logarithm of odds (LOD) value, nearest marker to the QTL peak, P-value of the QTL peak, percent phenotypic variance exmplained (% var R^2^), and additive effect size.

| Year | QTL | Chr. | Peak Pos.<br>(cM) | LOD<br>value | Nearest<br>marker | P-value | % var<br>($R^2$) | Additive<br>effect |
| --- | --- | --- | --- | --- | --- | --- | --- | --- |
| 2019 | *q19-1* | 4L | 572.9 | 4.56 | *umc2139* | 0.007 | 8.62 | -0.422 |
|  | *q19-2* | 5S | 248 | 3.88 | *umc1447* | 0.029 | 7.4 | -0.38 |
| 2020 | *q20-1* | 4L | 563 | 4.31 | *AY110170* | 0.009 | 9.09 | -0.32 |
|  | *q20-2* | 5S | 245 | 4.09 | *umc1557* | 0.025 | 8.53 | -0.3 |

### QTL on chromosome 4

In 2019, we identified a strong QTL (q19-1) on chromosome 4L that has a LOD score of 4.56 (*p =* 0.007) **(Figure 3, Table 1)**. Marker *p-umc2139* was the nearest peak marker located at 572.9 cM on the genetic map. The maker showed a negative additive effect (−0.422) and explained 8.62% of total phenotypic variation **(Table 1)**. We calculated the confidence interval of the significant QTL using a 1.5 LOD interval with the nearest marker **(Table 2)**. We found marker *p-umc2188* around 554.1 cM (constructed on *umc2188* gene that starts from 200,113,752 bp; bin 4.08) as the lower flanking marker while *p-php10025* around 580.0 cM (nearest gene *cbp2*, at 235,806,854 bp; bin 4.09) as the upper flanking marker to the peak marker *p-umc2139* **(Table 2)**. Based on 1.5 LOD confidence interval, q19-1 was narrowed down to a 25.9 cM interval size, or 35.7 Mb region based on B73 RefGen_v3.

**Table 2:**
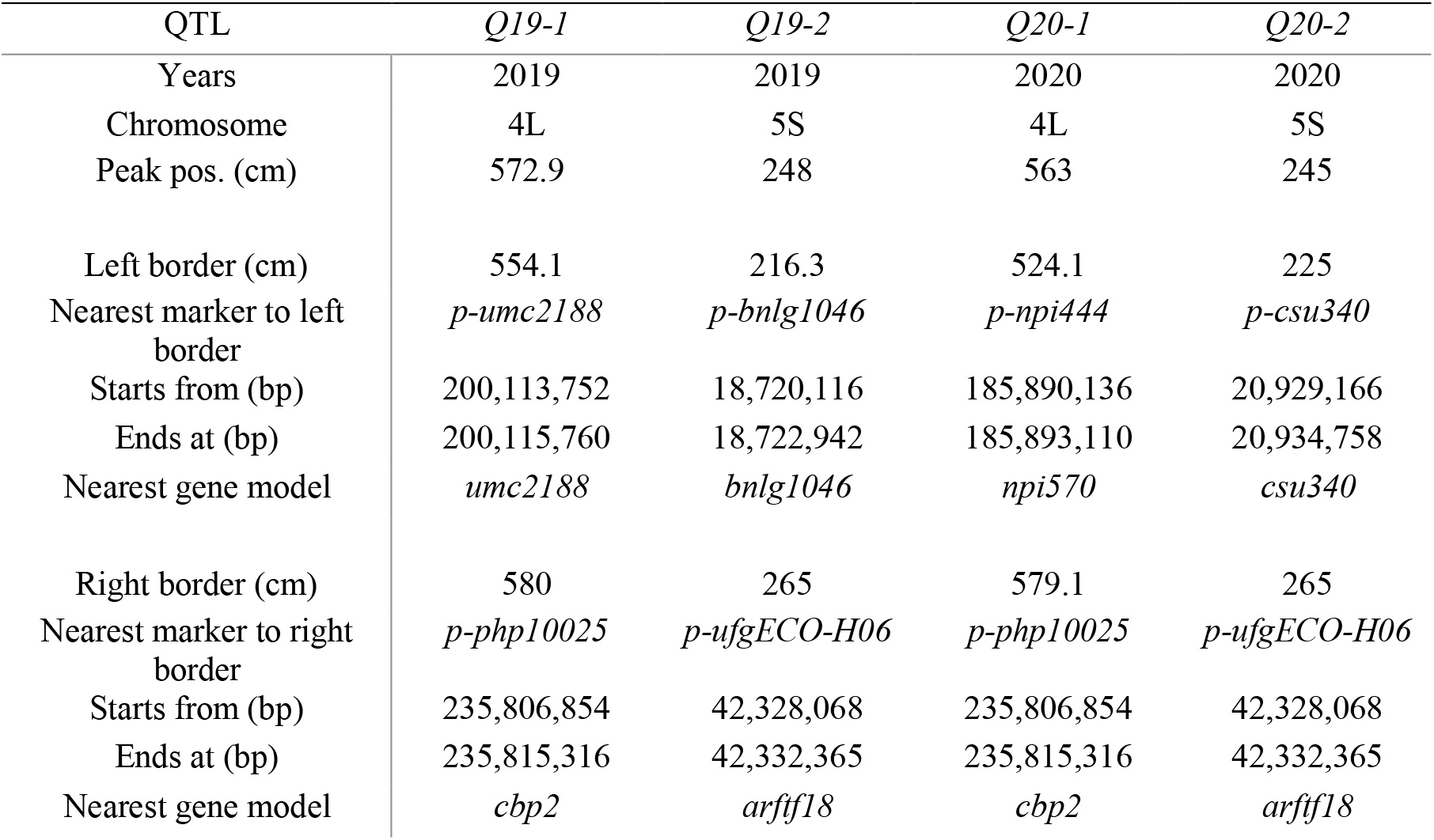
List of significant QTL with their nearest markers, confidence interval of 1.5 LOD support interval with the nearest marker, physical position (in bp) on genetic map for 2019 and 2020. The physical positions of marker were based on B73_RefGen v3.

| QTL | *Q19-1* | *Q19-2* | *Q20-1* | *Q20-2* |
| --- | --- | --- | --- | --- |
| Years | 2019 | 2019 | 2020 | 2020 |
| Chromosome | 4L | 5S | 4L | 5S |
| Peak pos. (cm) | 572.9 | 248 | 563 | 245 |
| Left border (cm) | 554.1 | 216.3 | 524.1 | 225 |
| Nearest marker to left border | *p-umc2188* | *p-bnlg1046* | *p-npi444* | *p-csu340* |
| Starts from (bp) | 200,113,752 | 18,720,116 | 185,890,136 | 20,929,166 |
| Ends at (bp) | 200,115,760 | 18,722,942 | 185,893,110 | 20,934,758 |
| Nearest gene model | *umc2188* | *bnlg1046* | *npi570* | *csu340* |
| Right border (cm) | 580 | 265 | 579.1 | 265 |
| Nearest marker to right border | *p-php10025* | *p-ufgECO-H06* | *p-php10025* | *p-ufgECO-H06* |
| Starts from (bp) | 235,806,854 | 42,328,068 | 235,806,854 | 42,328,068 |
| Ends at (bp) | 235,815,316 | 42,332,365 | 235,815,316 | 42,332,365 |
| Nearest gene model | *cbp2* | *arftf18* | *cbp2* | *arftf18* |

Similarly, in 2020, we identified QTL (q20-1) on chromosome 4L, which has a LOD score of 4.31 and (*p* = 0.009) **(Figure 3, Table 1)**. Marker *AY110170* was near the peak and was located at 563 cM on the genetic map. The maker showed a negative additive effect (−0.32) and explained 9.09% of total phenotypic variation **(Table 1)**. Using a 1.5 support interval with the nearest marker, we found marker *p-npi444* around 524.1 cM (nearest gene *npi570* at Chr4: 185,488,773 bp; bin 4.08) as the lower flanking marker while *p-php10025* around 580.0 cM (nearest gene *cbp2*, at 235,806,854 bp; bin 4.09) as the upper flanking marker to the peak marker *AY110170* **(Table 2)**. Based on a 1.5 LOD confidence interval, q20-1 is narrowed down to a 55.9 cM interval size or to a 49.9 Mb region based on B73 RefGen_v3, which was 14.2 Mb wider than that observed in 2019.

### QTL on chromosome 5

In 2019, we identified another QTL (q19-2) on chromosome 5S that has a LOD score of 3.88 and (*p* = 0.029) **(Figure 3, Table 1)**. Marker *p-umc1447* was the nearest peak marker located at 248 cM on the genetic map. The maker showed a negative additive effect (−0.38) and explained 7.4% of total phenotypic variation **(Table 1)**. Using a 1.5 LOD confidence interval of the most significant QTL, we found marker *p-bnlg1046* around 216.3 cM (nearest gene *bnlg1046* at Chr5:18,770,116 bp; bin 5.03) as the lower flanking marker while *p-ufgECO-H06* around 265.0 cM (nearest gene *arftf18* at Chr5:42,326,568 bp; bin 5.03) as the upper flanking marker to the peak marker *p-umc1447* **(Table 2)**. Based on a 1.5 LOD confidence interval, q19-2 is narrowed down to a 23.6 Mb region based on B73 RefGen_v3.

In 2020, we identified the same QTL (q20-2) as the 2019 data on chromosome 5S, which has a LOD score of 4.09 and a *p-value* of 0.025 **(Table 1)**. We found marker *p-umc1557* as the nearest peak marker located at 245 cM in the genetic map. The maker showed a negative additive effect (−0.3) and explained 8.53 % of total phenotypic variation **(Table 1)**. Using a 1.5 support interval with the nearest marker, we found marker *p-csu340* around 225 cM (nearest gene *csu340* at Chr5:20,929,166 bp; bin 5.03) as the lower flanking marker while *p-ufgECO-H06* around 265.0 cM (nearest gene *arftf18* at Chr5:42,326,568 bp; bin 5.03) again as the upper flanking marker to the peak marker *p-umc1557* **(Table 2)**. Based on a 1.5 LOD confidence interval, q20-2 is narrowed down to a 21.4 Mb region based on B73 RefGen_v3.

### Allele from Mo17 in conjunction with *Tcb1-s* work more effectively to block the foreign pollen when compared to the B73 allele in with *Tcb1-s*

We were interested in understanding marker effects from the two of the founder lines, B73 with *Tcb-s1* and Mo17 with *Tcb1-s*, in effectively blocking *tcb1* pollen. We used the peak markers from our significant QTL on 4L and 5S from both years and associated it with the mean contamination score. Both peak markers from QTL 4L in 2019 (p-umc 2139) and 2020 (AY110179) were found to work more effectively in a Mo17/*Tcb1-s* genetic background in blocking the foreign pollen (*tcb1*), and worked less effectively in a B73/*Tcb1-s* genetic background in blocking the foreign pollen **(Figure 4A,C)**. Similarly, both peak markers from QTL 5S in 2019 (p-umc 1447) and 2020 (p-umc1557) were found to work more effectively in the Mo17/*Tcb1-s* genetic background in blocking the foreign pollen (tcb1) than the B73/*Tcb1-s* genetic background **(Figure 4B,D)**.

**Figure 4:**
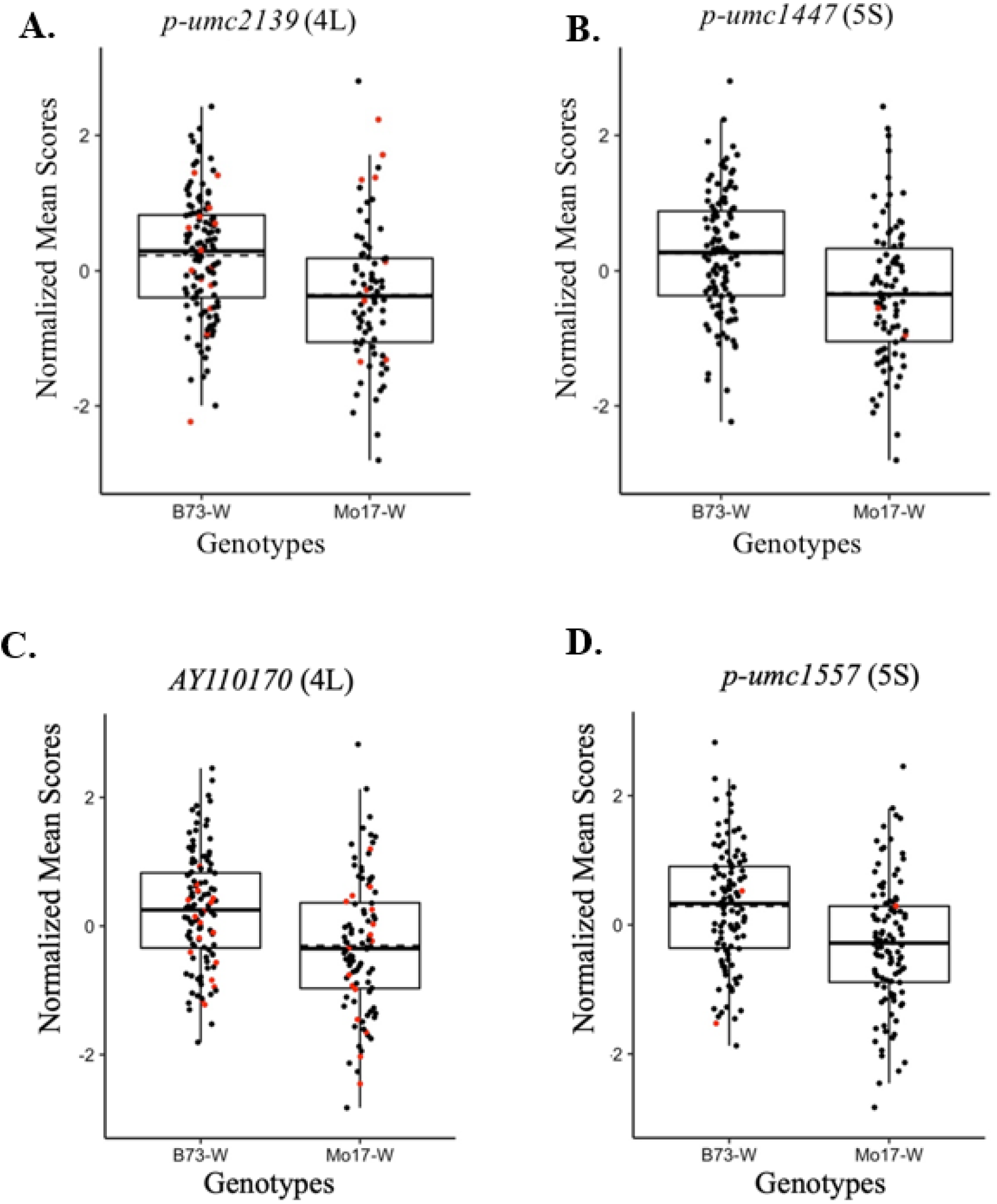
Peak markers effect on the mean contamination score of *Tcb1* from QTL analysis in 2019 and 2020. **A)** Phenotype X Genotype (PXG) plot for a significant marker (*p-umc2139*) on 4L from 2019. **B)** PXG plot for a significant marker (*p-umc1447*) on 5S from 2019. **C)** PXG plot for a significant marker (*AY110170*) on 4L from 2020. **D)** PXG plot for a significant marker (*p-umc1557*) on 5S from 2020. Boxplots showing phenotypic scores of significant markers with respective B73 and Mo17 allele. X-axis represents alleles and Y-axis represents normalized mean score for each line. The dots indicate the actual mean value for each observation. Red dots mean phenotype imputed values. Black dash line indicates mean contamination score while solid black line indicates median value.

### Candidate genes in the stable QTL regions of 4L and 5S

We were then interested in identifying potential candidate genes within the stable QTL regions that overlapped in 2019 and 2020 for both 4L and 5S. For the QTL on 4L, we found that the intervals from 2019 and 2020 overlapped by 35.7 Mb **(Figure 5A)**. We prioritized this overlapped region to study candidate genes that could potentially modify the efficacy of *Tcb1-s*. This overlapping region of chromosome 4 harbors 47 protein-coding genes **(Figure 5A, Supplemental Table 3)**. Among them were nine candidate genes possibly affecting the pollen blocking efficacy of *Tcb1-s* **(Table 3)**. For QTL on 5S, the QTL interval from 2019 was 2.2 Mb narrower than that observed in 2020 **(Figure 5B)**. The overlapping region on chromosome 5S harbors 45 protein-coding genes **(Figure 5B, Supplemental Table S4)**. Among them, we found three promising candidate genes that might have contributed to the *Tcb1-s* gene activity **(Table 3)**.

**Figure 5:**
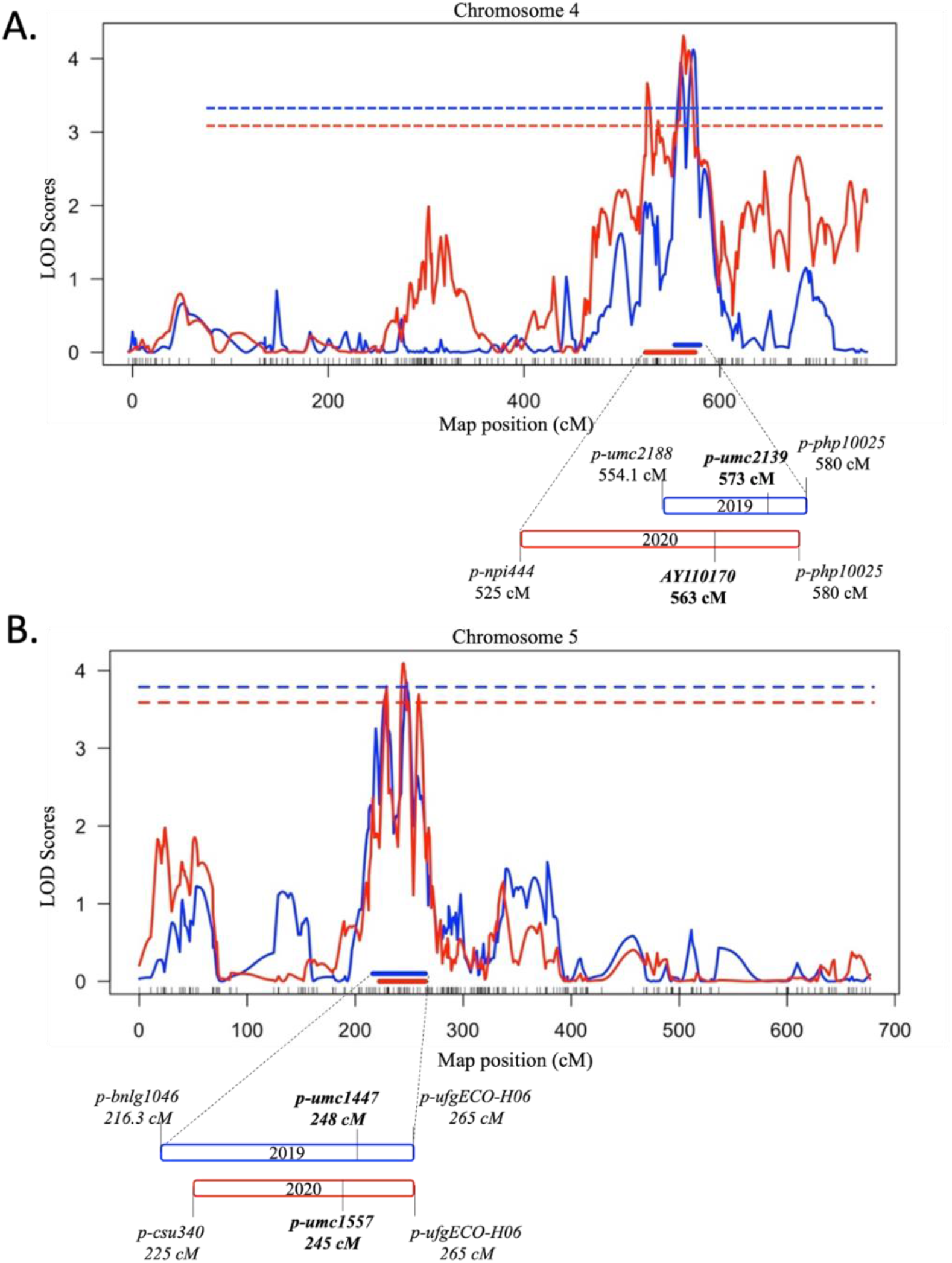
Stable QTL overlapping regions of 4L and 5L in 2019 and 2020. **A)** Overlapping QTL region on 4L; the blue and red line graph indicates QTL map from 2019 and 2020, respectively. Similarly, horizontal blue and red dotted line indicates 1000 permutated LOD threshold value of 3.70 (2019) and 3.66 (2020) respectively. It also shows the peak markers and the flanking markers in 2019 and 2020 QTL along with the overlapping region. **B)** Overlapping QTL region on 5S; the blue and line graph indicates QTL map from 2019 and 2020, respectively. Similarly, horizontal blue and red dotted line indicates 1000 permutated LOD threshold value of 3.70 (2019) and 3.66 (2020), respectively. It also shows the peak markers and the flanking markers in 2019 and 2020 along with its overlapping region.

**Table 3:** Candidate genes on 4L and 5L

| QTL | Gene name | Gene annotation |
| --- | --- | --- |
| 4L | GRMZM2G465165 | Putative lectin-domain receptor-like protein kinase family protein |
| 4L | GRMZM2G114471 | Chalcone synthase 8 |
| 4L | GRMZM2G125206 | F-box domain containing protein |
| 4L | GRMZM2G027336 | Glyoxal oxidase |
| 4L | GRMZM2G035843 | Putative calcium-dependent protein kinase family protein |
| 4L | GRMZM2G060800 | Aldehyde dehydrogenase |
| 4L | GRMZM2G016558 | Putative ankyrin repeat domain family protein |
| 4L | GRMZM2G048353 | Defensin |
| 4L | GRMZM2G437040 | Defensin |
| 5S | GRMZM2G018082 | Oxidoreductase |
| 5S | GRMZM2G087827 | Esterase |
| 5S | GRMZM2G094599 | Armadillo |

## Discussion

In this study, we introgressed *Tcb1-s* into the IBM RILs to map the modifiers of the *Tcb1*. A detailed understanding of these modifiers is important to understand the action of the *Tcb1* genetic system and to study the range of pollen blocking effects via different combinations of modifiers. Previously, eight modifiers (LOD > 2) for *Tcb1-s* were identified using 77 out of 302 IBM RILs (Burch 2018).

Limited sample size and small LOD values may have contributed to the small QTL obtained. In this study, we used a larger sample size to map QTL and identified two significant QTL on 4L and 5S consistent between years. A similar study searching for the modifiers of the gametophyte factor *Ga1-s* showed evidence of significant QTL on chromosomes 4 and 10 (Shrestha 2016). Similarly, a genome-wide association study by Hurst (2019) identified seven significant SNPs as possible modifiers of the *Ga1-s* system (Hurst et al. 2019). It is currently unknown whether these modifiers and their molecular mechanisms are unique or shared between cross-incompatibility systems.

We mapped QTL exhibiting negative additive effects contributed by the Mo17 allele that contributed to blocking foreign pollen (**Table 1**). This negative effect was supported by the percent mean color contamination of the F1s of (B73 x *Tcb1-s*) and (Mo17 x *Tcb1-s*) in that the mean color contamination was higher in the (B73 x *Tcb1-s)* and lower in the (Mo17 x *Tcb1-s)* for both years **(Supplemental Figure 1)**. Although we have demonstrated that the mean allelic effect from Mo17 in conjunction with *Tcb1-s* works more effectively to block the foreign pollen effectively when compared to B73 with *Tcb1-s*, we want to highlight that we observed phenotypic differences (variable color contamination scores) among the different ears of the same genetic background **(Supplemental Figure 1)**. One potential explanation for such observation is that we did not consider the silks’ age among ears of the same genetic background during the pollination. Future experiments should consider silk age during pollination and try to control for this variability during pollination to avoid such discrepancies. In addition, we also observed a lower amount of blue-colored (*tcb1*) kernel contamination in 2020 compared with 2019 **(Supplemental Figure 1)**. This could be due to several reasons; (1) the use of two different color markers for *tcb1* pollen across years could possibly affect the blocking activity of *Tcb1-s*, (2) the age of the silks could impact *Tcb1-s* activity, and (3) several other environmental factors during the growth years may affect the *Tcb1-s* efficacy.

Using two years of replicated field trials, we identified a QTL on chromosome 4L that we narrowed down to 35.7 Mb **(Figure 5A)** that contained 47 candidate genes **(Figure 5A, Supplemental Table 3)**. Nine out of 47 genes were related to the pollen-pistil interaction and incompatibility **(Table 3**.

Among them, *GRMZM2G035843* is annotated as a putative calcium-dependent protein kinase. In *Nicotiana tabacum*, calcium-dependent protein kinase (CPK32) activates a Ca^2+^ permeable channel in the plasma membrane of pollen tube tips, which increases the influx of Ca^2+^, resulting in the polar growth of pollen tubes (Zhou et al. 2014). Ca^2+^ dependency is observed from pollen germination to the end of pollen tube elongation. During the pectin methyl de-esterification in the pollen tube, the resulting carboxyl groups are then bound by Ca^2+^ ions, forming cross-linking among pectin molecules. De-esterification produces a rigid cell wall, whereas esterification softens the cell wall. This suggests that the concentration of Ca^2+^ directly affects the cell wall elongation of a pollen tube (Hepler 2005). Therefore, our identified calcium-dependent protein kinase may be a possible modifier of *Tcb1-s*.

Another candidate gene, *GRMZM2G016558*, is annotated as a putative ankyrin repeat domain family protein which is known to be involved in the polar growth of pollen tubes. Ankyrin repeat domain proteins (LlANK) have been shown to contribute to the polar growth of pollen tubes in lily (Huang et al. 2006). Two additional candidate genes on 4L, *GRMZM2G048353* and *GRMZM2G437040*, were annotated as *Defensin* proteins. A related defensin-like protein ZmES4, was found to induce interaction with potassium in pollen tube tips in maize (Amien et al. 2010). In *Arabidopsis*, the defensin-like gene (*AtLURE1*) encodes an attractant for cell-to-cell communication between the pollen tube and pistil. This protein is produced in the synergid and is diffused through the micropyle. It helps to guide the protruding pollen tubes towards the ovule. Gametophytic mutants with defective micropylar guidance showed low expression of *AtLURE1*, resulting in impaired pollen tube guidance towards the ovary (Takeuchi and Higashiyama 2012). In incompatible pollinations of *tcb1* pollen onto *Tcb1-s* silks, this gene might be expressed at a low level, resulting in a lack of pollen tube guidance and eventual blockage by *Tcb1-s* activity.

*GRMZM2G1255206* encodes an F-box domain containing protein. F-Box proteins affect pollen tube growth inhibition in incompatible pollinations through ubiquitin-mediated protein degradation (Kumar and McClure 2010). In *Antirrhinum hispanicum*, AhSLFS2 is an F-Box protein that mediates the activation of a selective yet destructive, S-RNase during self-incompatible pollinations responses (Qiao et al. 2004). Another candidate gene *GRMZM2G060800* encodes an aldehyde dehydrogenase necessary to biosynthesize indole-3-acetic acid (IAA). A similar candidate plant hormone gene, *GRMZM2G089856*, encodes for a 1-aminocyclopropane-1-carboxylate oxidase necessary to biosynthesize ethylene. Different studies have found these hormones to be associated with pollination compatibility. Auxin production was increased in compatible pollination (Hasenstein and Zavada 2001), whereas incompatible pollination induced ethylene production in pistil tissues resulting in incompatible pollination. In self-incompatible species of *Petunia inflata*, ethylene production was triggered in pistils during the early stage of pollen tube growth (Holden et al. 2003).

*GRMZM2G027336* encoding glyoxal oxidase is known to be a putative cell wall expansion protein that plays a key role in cell wall metabolism. This protein is highly expressed in fertile flowering buds, whereas the expression was low in flowering buds of cotton with cytoplasmic male sterility (SUZUKI ET AL. 2013). *GRMZM2G114471* encoding chalcone synthase catalyzes the biosynthesis of flavonoids (a signal for successful pollen tube growth). A chalcone synthase mutant fails to produce functional pollen tubes in maize and petunia (Mo et al. 1992). Therefore, these genes can be considered important candidate genes underlying the QTL at chromosome 4 that affect *Tcb1* blocking action.

The QTL found in both years on chromosome 5 was narrowed down to a 24.1Mb region **(Figure 5B)** that consists of 45 candidate genes **(Figure 5B, Supplemental Table 4)**. Three out of forty-five genes were related to pollen-pistil interactions and incompatibility **(Table 3)**. One promising gene which might affect the efficacy of pollen blockage is *GRMZM2G087827*. This gene encodes an esterase, an important enzyme family in plant reproduction. Different forms of esterase are necessary for successful fertilization in plants. In a dry-type stigma, pollen dissolves the cutin layer of the stigma through the activity of carboxyl esterases present in pollen (Rejón et al. 2012). Pollen tube growth is regulated inside pistils by different classes of pectin methylesterases (PME). ZmPME10-1 is predominantly expressed at the apex of growing pollen tubes and helps maintain pectin methylesterification necessary for pollen tube growth in *Ga1-s* silks (Zhang et al. 2018). Other PME proteins like ZmPME3 (encoded by *Ga1-s)* (Moran Lauter et al. 2017) and TCB1-F/PME38 (encoded by *Tcb1*-s) (Lu, Hokin, Kermicle, Hartwig and Evans 2019) are pistil expressed and play an important role in successful fertilization. Therefore, *GRMZM2G087827* might have similar actions to the esterases above, making it a strong candidate modifier of *Tcb1-s* activity.

The QTL region on 5S contains another gene, *GRMZM2G018082*, annotated as an oxidoreductase. This protein has been found to affect self-incompatibility in different crops. An A-like oxidoreductase has been identified as a candidate gene causing pollen rejection due to redox regulation in apricot (Muñoz-Sanz et al. 2017). Similarly, in *Solanum habrochaites*, an NAD(P)- linked oxidoreductase and lipid oxidoreductase are known to be involved in a redox reaction resulting in unilateral incompatibility (Broz *et al*. 2017). Lastly, *GRMZM2G094599* was annotated as an armadillo/beta-catenin-like repeat protein. ARMADILLO REPEAT ONLY1 (ARO1) showed functions of F-actin organization in the pollen tube tip and polar growth in *Arabidopsis thaliana* (Gebert et al. 2008). Based on their known putative functions reported in other species, these genes could be important candidates associated with the QTL on chromosome 5.

## Conclusion

We have identified two stable QTL on chromosomes 4L and 5S that modify *Tcb1-s* action in the IBM population. We show that the allele from Mo17, in conjunction with *Tcb1-s*, works more effectively to block foreign *tcb1* pollen when compared to the B73 allele. Finally, using data from multiple years, we narrowed down the genomic interval of *Tcb1-s* modifiers to identify 12 candidate genes that have direct or indirect effects on the cell wall pectin esterification and de-esterification balance, polarized pollen tube growth, cell wall expansion, and pollen-pistil interactions. All candidate effects could potentially lead to the observed *Tcb1-s* cross incompatibility barrier. Our findings provide exciting possibilities for the mechanistic understanding of *Tcb1-s* action under different genetic backgrounds and will further aid in the marker development and introgression of *Tcb1* in genetic backgrounds that require controlled pollination.

## Data Availability Statement

Data for this publication are available in the supplementary information.

## Author Contributions

NM and MKB performed the experiments, processed and analyzed the data and wrote the manuscript with VS. PA assisted in field experiments and data collection and revisions to the manuscript. AG provided scientific output and assisted in writing the manuscript. NM, MKB, VS, and DLA designed the experiments, participated in result interpretation, and assisted with writing and revisions to the manuscript. YW supervised the work, provided scientific output and critically revised the manuscript. DLA conceived the study, supervised the work and its coordination, acquired funding, and assisted with interpretation of results and revisions to the manuscript. All authors contributed to the text and approved the final manuscript.

## Acknowledgments

We would like to dedicate this work to our late advisor Dr. Donald Leon Auger and our late collegue Abiskar Gyawali. We thank to the South Dakota State University Agricultural Experiment Station for their assistance in field experiments.

