## Supplemental Figures 1-5 for "Genomic mapping of the modifiers of *teosinte crossing barrier 1* (*Tcb1*)"

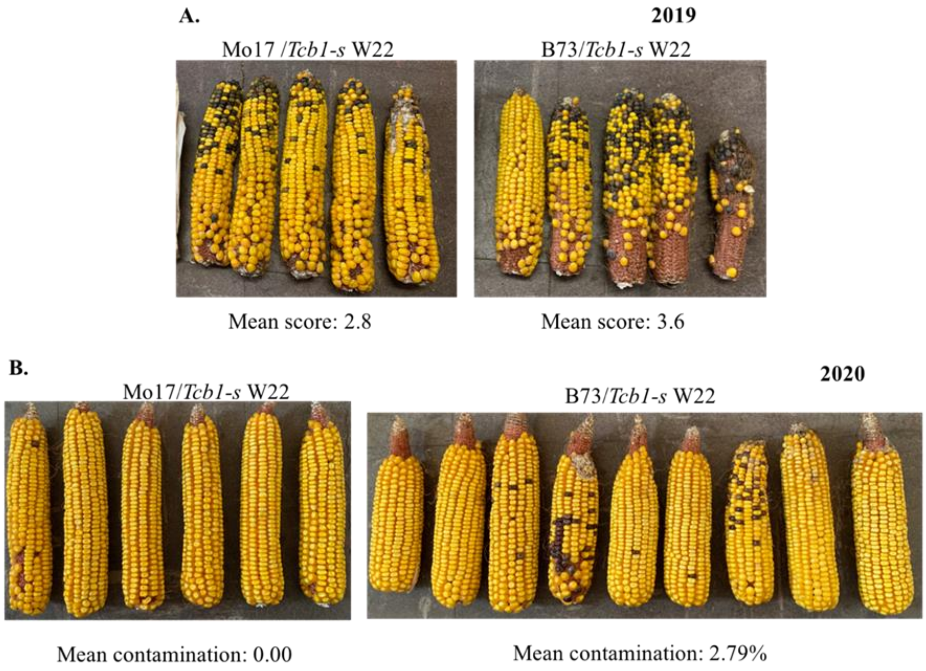


**Supplemental Figure 1:** Mean color kernel contamination in controls (Mo17/*Tcb1-s* W22 and B73/*Tcb1-s* W22) from 2019 and 2020. **A.** The color contamination in controls in 2019. The mean score shown was based on the standardized scale. **B.** The color contamination in controls in 2020. The mean contamination represented actual percentage of color kernels in the ears.

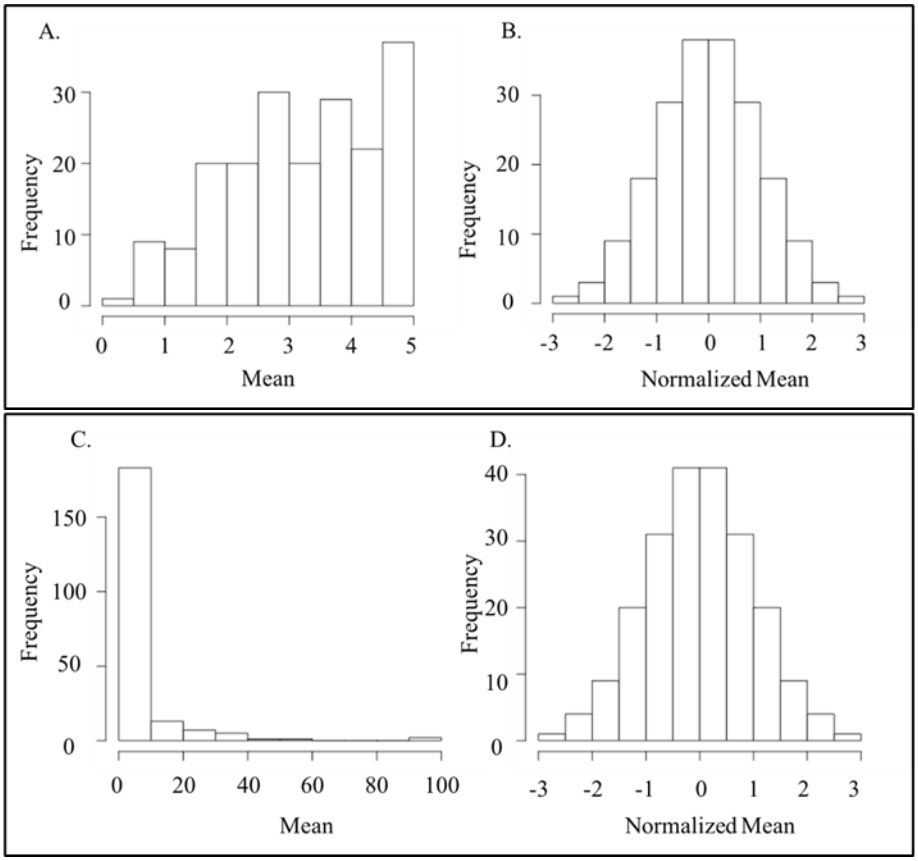


**Supplemental Figure 2**: Histogram showing the distribution of phenotypic data before and after normalization. The histogram for the phenotype date before quantile normalization is represented by A, C and E. B, D and F represent the data after normalization. A, B, C, and D show the phenotypic data from the QTL mapping population in 2019 and 2020.

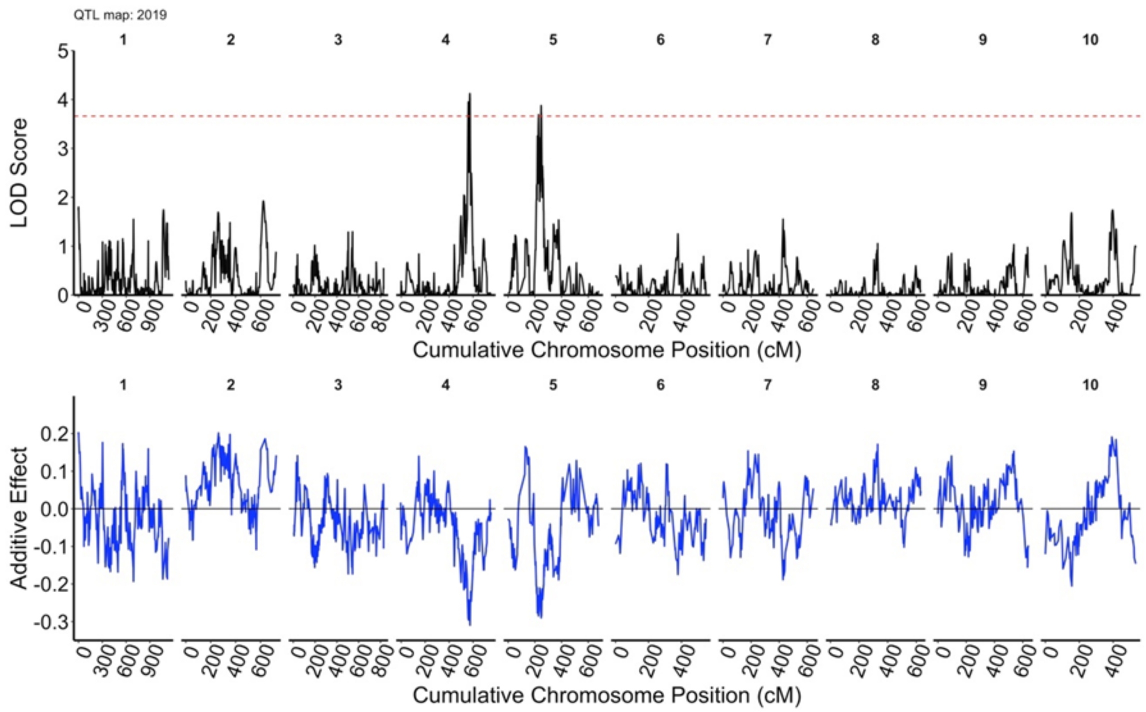


**Supplemental Figure 3**: QTL and genome wide effect scan for 2019. **A.** QTL map obtained by analyzing normalized values of phenotype scoring. The red dotted line indicates 1000 permutated LOD threshold value of 3.70. **B.** Estimated QTL additive effect along all chromosomes for the identified QTL.

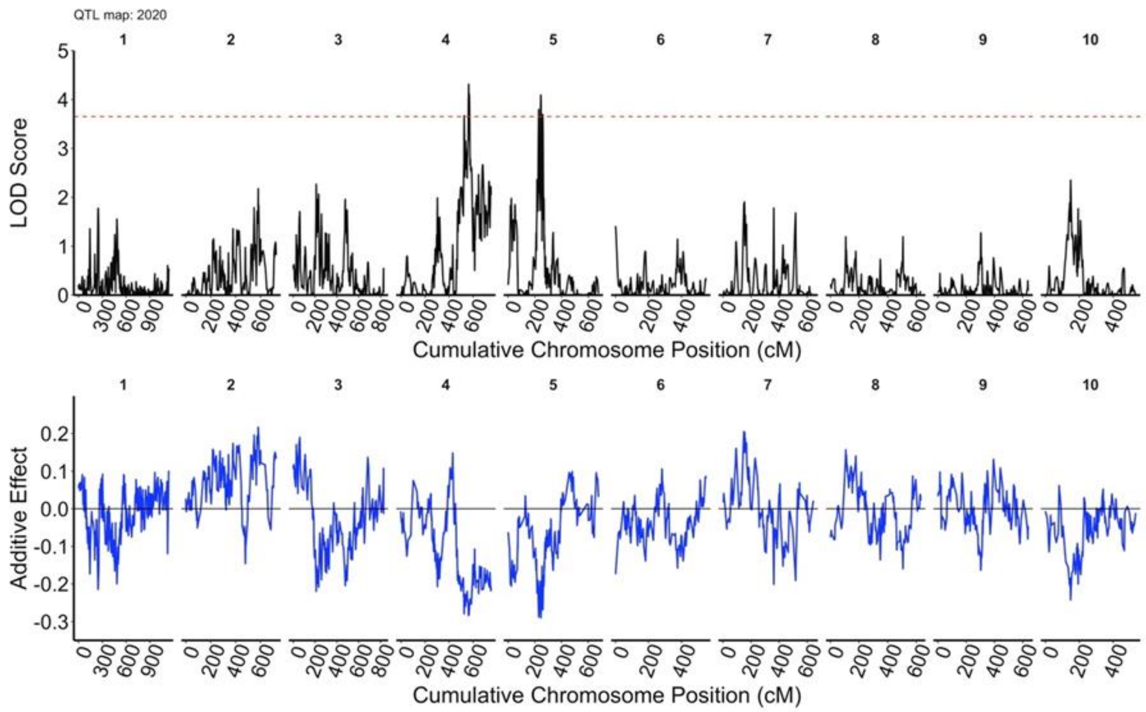


**Supplemental Figure 4**: QTL and genome wide effect scan for 2020. **A.** QTL map obtained by analyzing normalized values of phenotype scoring. The red dotted line indicates 1000 permutated LOD threshold value of 3.66. **B.** Estimated QTL additive effect along all chromosomes for the identified QTL.

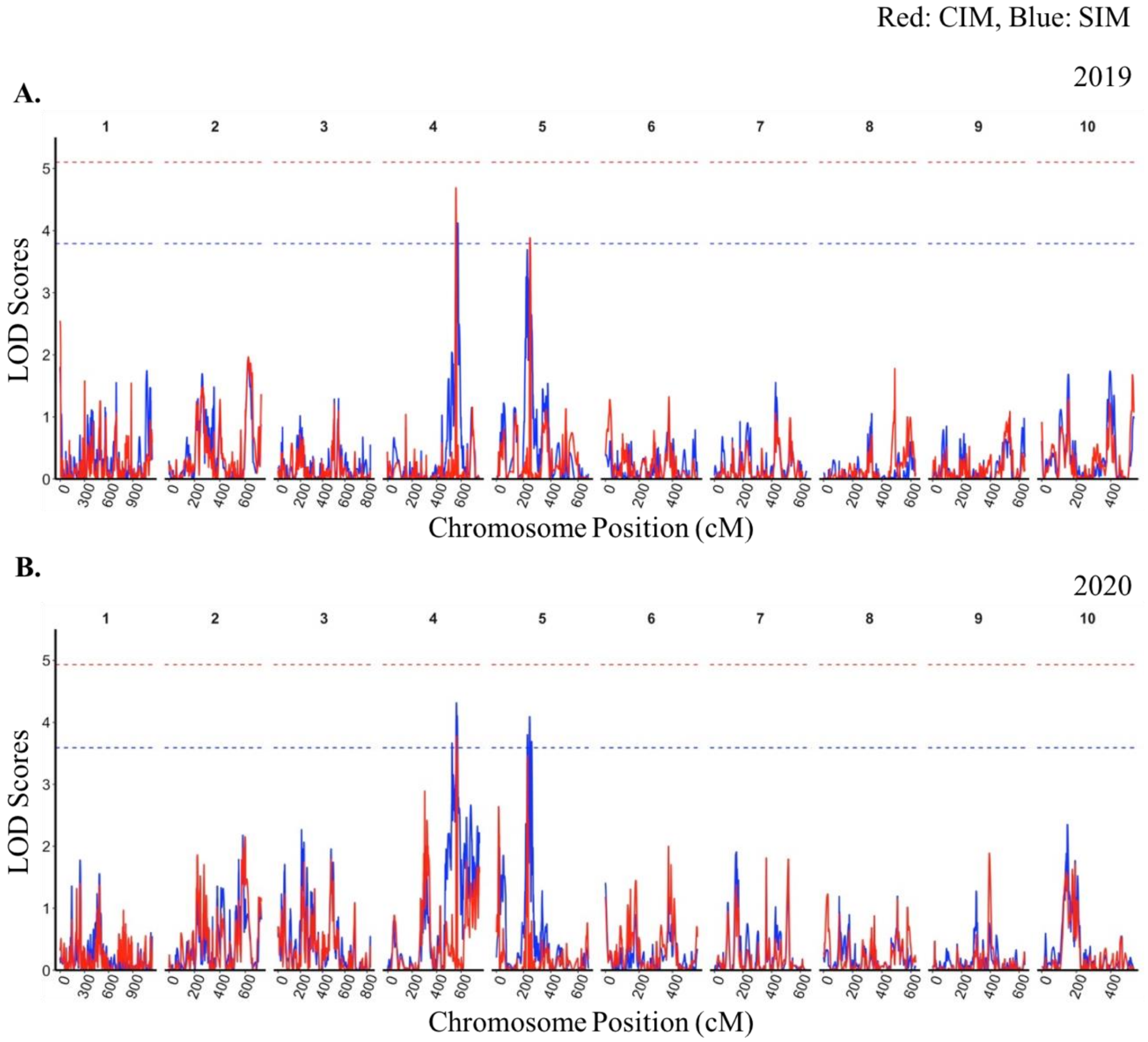


**Supplemental Figure 5:** Overlaying QTLs obtained from simple interval mapping (SIM) and composite interval mapping (CIM) **A.** Overlaying QTL maps experiment in 2019. The vertical blue line indicates the QTLs using SIM while the vertical red line indicates QTLs from CIM. Similarly, the horizontal blue and red dotted line indicates 1000 permutated LOD threshold value of 3.70 (SIM) and 4.90 (CIM), respectively. **B.** Overlaying QTL maps of experiment in 2020. The vertical blue line indicates the QTLs using SIM while the vertical red line indicates QTLs from CIM. Similarly, the horizontal blue and red dotted line indicates 1000 permutated LOD threshold value of 3.66 (SIM) and 4.90 (CIM), respectively.
